## Supplementary for "Novel Variance-Component TWAS method for studying complex human diseases with applications to Alzheimer’s dementia"

### Supplemental Texts

#### Text S1. Details of PrediXcan's approach of estimating cis-eQTL effect sizes

PrediXcan TWAS method [1] employs Elastic-Net penalized regression method [2] to estimate cis-eQTL effect sizes $\boldsymbol{w}$ from Equation (1) in the main text. Basically, the Elastic-Net method assumes a combined LASSO ($L_1$) [3] and Ridge ($L_2$) [4] penalty and estimate $\boldsymbol{w}$ by the following equation

$$\widehat{\boldsymbol{w}} = \underset{\boldsymbol{w}}{argmin}(\left\|\boldsymbol{E_g} - \boldsymbol{Gw}\right\|_2^2 + \lambda(\alpha||\boldsymbol{w}||_1 + \frac{1}{2}(1-\alpha)||\boldsymbol{w}||_2^2)) ,$$

where $\|\cdot\|_1$ denotes $L_1$ norm, $\|\cdot\|_2$ denotes $L_2$ norm. Particularly, $\alpha$ is taken as 0.5 by PrediXcan method [1] and penalty parameter $\lambda$ can be tuned by a 5-fold cross validation.

#### Text S2. Details of TIGAR's approach of estimating cis-eQTL effect sizes

TIGAR[8] provides a more flexible approach to nonparametrically estimate cis-eQTL effect sizes $\boldsymbol{w}$ from Equation (1) in the main text by a Bayesian DPR method [5]. The DPR method assumes a normal prior distribution $N(0, \sigma_w^2)$ for cis-eQTL effect sizes and a Dirichlet process prior [6] for effect-size variance $\boldsymbol{\sigma_w^2}$ as follows:

$$\mathrm{w_i} \sim \mathrm{N}(0, \sigma_\mathrm{w}^2), \qquad \sigma_\mathrm{w}^2 \sim \mathrm{D}, \qquad \mathrm{D} \sim \mathrm{DP}(\mathrm{IG}(a, b), \xi).$$

That is, the prior distribution $D$ of effect-size variance deviates from a Dirichlet Process (DP) with an inverse gamma (*IG*) distribution and concentration parameter $\xi$. As proposed by previous studies, variational Bayesian algorithm [7, 8] is implemented to efficiently obtain posterior estimates $\widehat{\boldsymbol{w}}$.

**Text S3. Details of VC-TWAS approach with summary-level GWAS data and P-value calculation**

VC-TWAS with summary-level GWAS data

Since summary-level GWAS data are generally generated by meta-analysis based on the following single variant (SNP) regression model:

$$\boldsymbol{Y} = \boldsymbol{G}_{.j}\beta_j + \boldsymbol{\varepsilon}, \quad \varepsilon_i \sim N(0, \sigma_\epsilon^2) .$$

Here, the phenotype is assumed to be adjusted for other confounding covariates with mean 0, and the genotype vectors ($\boldsymbol{G}_{.j}$, j=1,…,m) are also assumed to be centered with mean 0. Without loss of generality, we assume GWAS summary statistics include the single variant effect size estimate $\hat{\beta}_j$ and corresponding standard error $\hat{\sigma}_j$ for the $j^{th}$ SNP, sample size $n$, and a reference LD covariance matrix $\boldsymbol{\Sigma}$ of all test SNPs.

Following the derivation provided by [9], given the marginal SNP effect size $\beta_j$ estimate $\widehat{\beta}_j = \frac{\boldsymbol{G}'_{.j}\boldsymbol{Y}}{\boldsymbol{G}'_{.j}\boldsymbol{G}_{.j}}$, the denominator $\boldsymbol{G}'_{.j}\boldsymbol{G}_{.j}$ can be approximated by using the $jth$ diagonal element of the reference LD covariance matrix, $\boldsymbol{\Sigma} \approx \boldsymbol{G}'\boldsymbol{G}/(n-1)$, with $\boldsymbol{G}'_{.j}\boldsymbol{G}_{.j} = (n-1)\Sigma_{j,j}$. Thus, the numerator of the score statistic for the $j^{th}$ SNP as shown in the main text (Equation (6)) can be estimated by

$$\boldsymbol{G}'_{.j}\boldsymbol{Y} = (n-1)\widehat{\beta}_j\Sigma_{j,j} .$$

In addition, based on the estimate for the marginal SNP effect size variance, the phenotype variance $\sigma_Y^2$ can be estimated by

$$\sigma_Y^2 = \frac{\boldsymbol{Y}'\boldsymbol{Y}}{(n-1)} \approx \frac{(\boldsymbol{G}'_{.j}\boldsymbol{G}_{.j})\widehat{\sigma_j^2}(n-1) + (\boldsymbol{G}'_{.j}\boldsymbol{G}_{.j})\widehat{\beta}_j^{\,2}}{(n-1)} = \Sigma_{j,j}\widehat{\sigma_j^2}(n-1) + \Sigma_{j,j}\hat{\beta}_j^{\,2}.$$

Since this estimate might vary with respect to the summary GWAS data of different SNPs, we take the median of $\Sigma_{j,j}\widehat{\sigma_j^2}(n-1) + \Sigma_{j,j}\hat{\beta}_j^{\,2}$ across all the SNPs as $\widehat{\sigma_Y}^2$ as suggested by the previous study [9].

Then the $Q$ statistic used by VC-TWAS using only GWAS summary-level data can be approximated by

$$Q = \sum_{j=1}^{m} w_j^2 \left( \frac{\boldsymbol{G'_{\cdot j} Y}}{\widehat{\sigma_Y}^2} \right)^2 .$$

P-value calculation for VC-TWAS

Under the null hypothesis, the $Q$ statistic used by VC-TWAS follows a mixture of chi-square distribution $\sum_{j=1}^{m} \lambda_j \chi_{j,1}^2$ [10, 11], where $(\lambda_1, \cdots, \lambda_m)$ are nonzero eigenvalues of $\boldsymbol{\Phi}$,

$$\boldsymbol{\Phi} = \boldsymbol{W \phi W}, \ \ \boldsymbol{\phi} = \boldsymbol{G' P G}, \boldsymbol{P} = \boldsymbol{V}^{-1} - \boldsymbol{V}^{-1} \boldsymbol{Z} \left( \boldsymbol{Z' V}^{-1} \boldsymbol{Z} \right)^{-1} \boldsymbol{Z' V}^{-1}$$

where $\boldsymbol{G}$ is the $n \times m$ genotype matrix, $\boldsymbol{Z}$ is the matrix of covariate data, $\boldsymbol{V} = \widehat{\sigma_Y}^2 \boldsymbol{I}$ for continuous traits with identify matrix $\boldsymbol{I}$, $\boldsymbol{V} = \boldsymbol{diag}[\boldsymbol{\hat{\mu}_1}(\boldsymbol{1} - \boldsymbol{\hat{\mu}_1}), \ldots, \boldsymbol{\hat{\mu}_n}(\boldsymbol{1} - \boldsymbol{\hat{\mu}_n})]$ for dichotomous traits.

If phenotype $\boldsymbol{Y}$ is centered and adjusted for other covariates as assumed when using summary-level GWAS data, then $\boldsymbol{\phi}$ can be simplified and approximated by $\boldsymbol{\phi} \approx \frac{(n-1)\boldsymbol{\Sigma}}{\widehat{\sigma_Y}^2}$ with reference LD covariance matrix $\boldsymbol{\Sigma}$ [12].

The p-value by VC-TWAS can then be conveniently obtained from several approximation and exact methods like the Davies exact method [13], which can be done by using both individual-level and summary-level GWAS data.

**Text S4. Details of ROS/MAP data**

In our applications of studying Alzheimer's dementia (AD), we used transcriptome and individual-level GWAS data generated for samples from the Religious Orders Study (ROS) and Rush Memory and Aging Project (MAP) [14-17] cohorts. ROS recruits nuns, priests, and brothers across the United States. MAP recruits participants living in private homes, subsidized housings, and retirement facilities across the greater Chicago metropolitan area. ROS/MAP data can be requested at www.radc.rush.edu.

Both studies employ harmonized data collection methods performed by the same staff for annual testing during life and for structured autopsy and collection of genomic data from blood and brain biospecimens. Harmonized data collection facilitates joint analyses of the studies' data. Details of the studies are described elsewhere.

We used microarray genotype data generated for 2,093 European-decent subjects from ROS/MAP [14-17], which are further imputed to the 1000 Genome Project Phase 3 [18]. Post-mortem brain samples (gray matter of the dorsolateral prefrontal cortex) from ~30% these ROS/MAP participants with assayed genotype data are also profiled for transcriptomic data by next-generation RNA-sequencing [19], which are used as reference data to train GReX prediction models in our application studies.

Using ROS/MAP data, we conducted TWAS for clinical diagnosis of late on-site Alzheimer's dementia (LOAD) as well as pathology indices of AD quantified with $\beta$-antibody and PHFtau specific immunostains. Quantitative pathology phenotypes $\beta$-amyloid load and PHFtau tangle density were studied. An additional phenotype of the summary measure of the burden of AD pathology (a combination of neuritic and diffuse plaques and neurofibrillary tangles based on modified Bielschowsky silver stain) [14, 15, 17] was also studied.

The tangle density quantifies the average PHFtau tangle density within two or more 20μm sections from eight brain regions — hippocampus, entorhinal cortex, midfrontal cortex, inferior temporal, angular gyrus, calcarine cortex, anterior cingulate cortex, and superior frontal cortex.

The $\beta$-amyloid load quantifies the average percent area of cortex occupied by $\beta$-amyloid protein in adjacent sections from the same eight brain regions. These two are based on immunohistochemistry. The global measure of AD pathology is based on counts of neuritic and diffuse plaques and neurofibrillary tangles (15 counts) on 6μm sections stained with modified Bielschowsky [14, 15, 17].

### Supplementary Figures

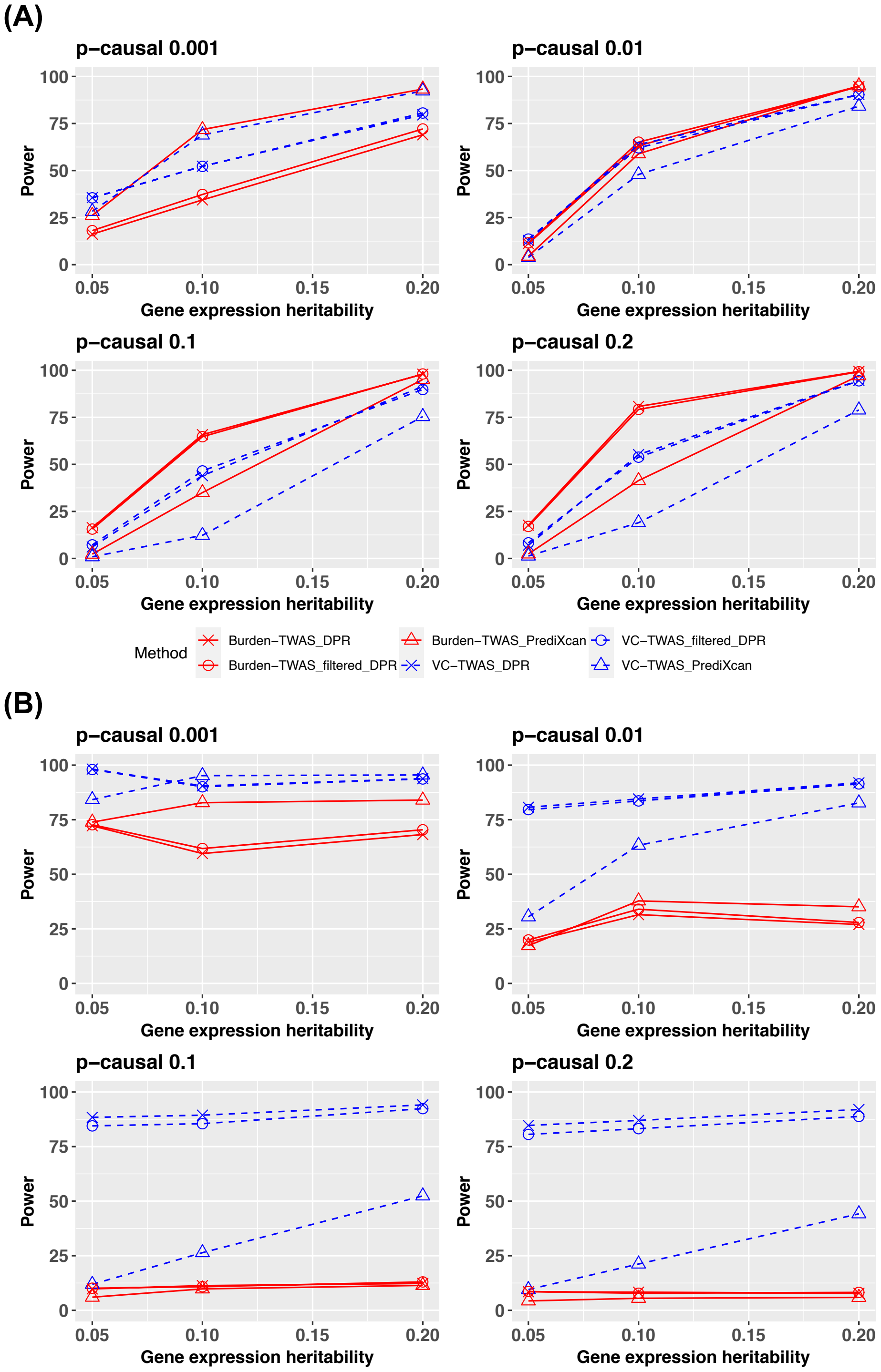


**Fig S1.** TWAS power comparison for VC-TWAS and Burden-TWAS with phenotypes simulated from Model I (A) and Model II (B). Various types of SNP weights were considered, including those derived from PrediXcan method, DPR method, and filtered DPR weights. In Model I, the combinations of causal probability and phenotype heritability are $(p_{causal}, h_p^2) = ((0.001,0.2), (0.01,0.3), (0.1,0.4), (0.2,0.5))$. In Model II, the combinations of causal probability and phenotype heritability are $(p_{causal}, h_p^2) = ((0.001,0.1), (0.01,0.1), (0.1,0.15), (0.2,0.15))$.

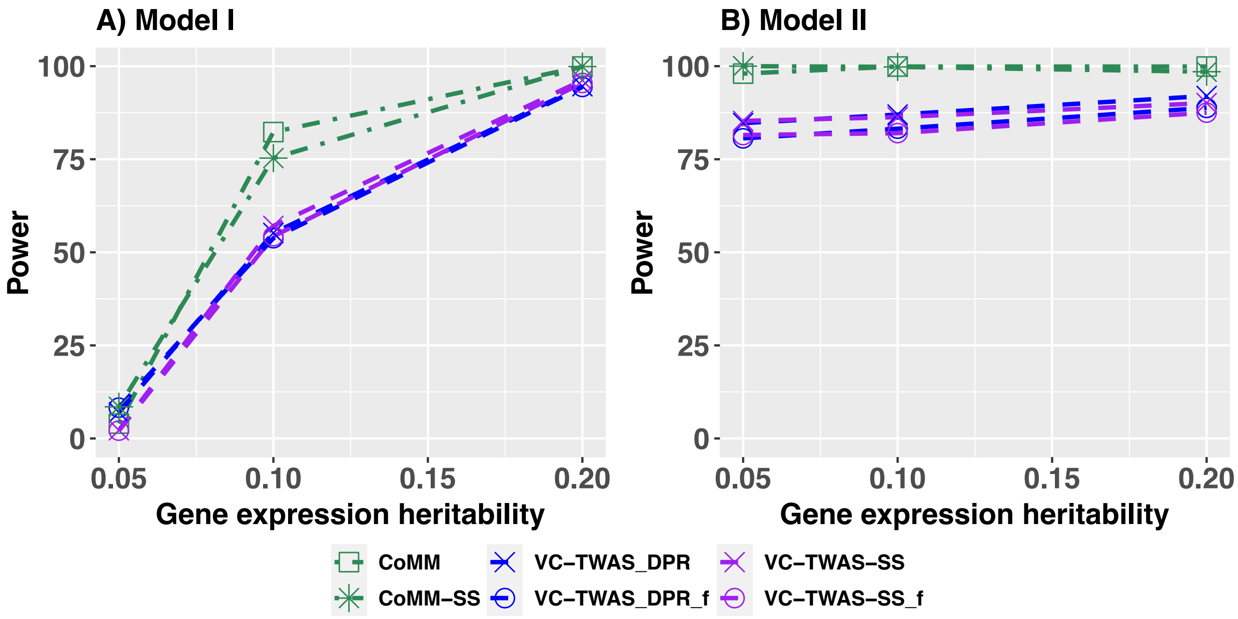


**Fig S2.** TWAS power comparison for VC-TWAS and CoMM with phenotypes simulated from Model I (A) and Model II (B) using individual-level and summary-level data under the scenarios with $p_{causal} = 0.2$.

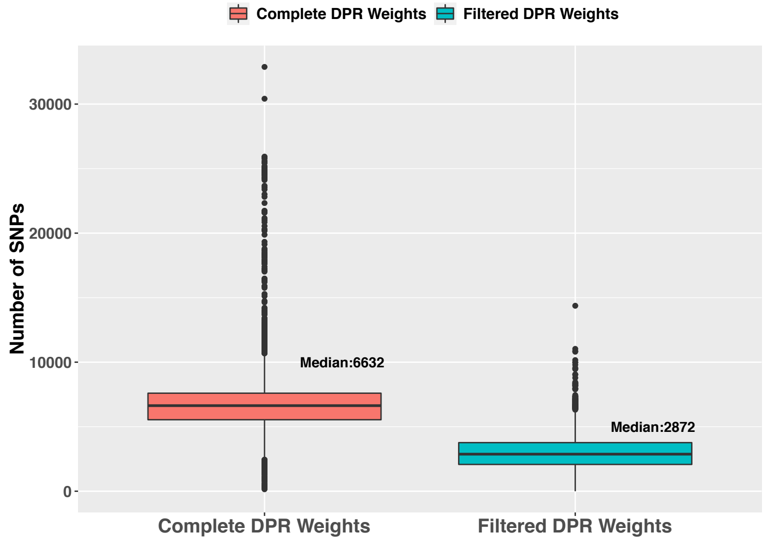


**Fig S3**. Box plot of the number of test SNPs considered by VC-TWAS of all genome-wide genes in the application studies of AD, with complete DPR weights and filtered DPR weights derived from the ROS/MAP training data.

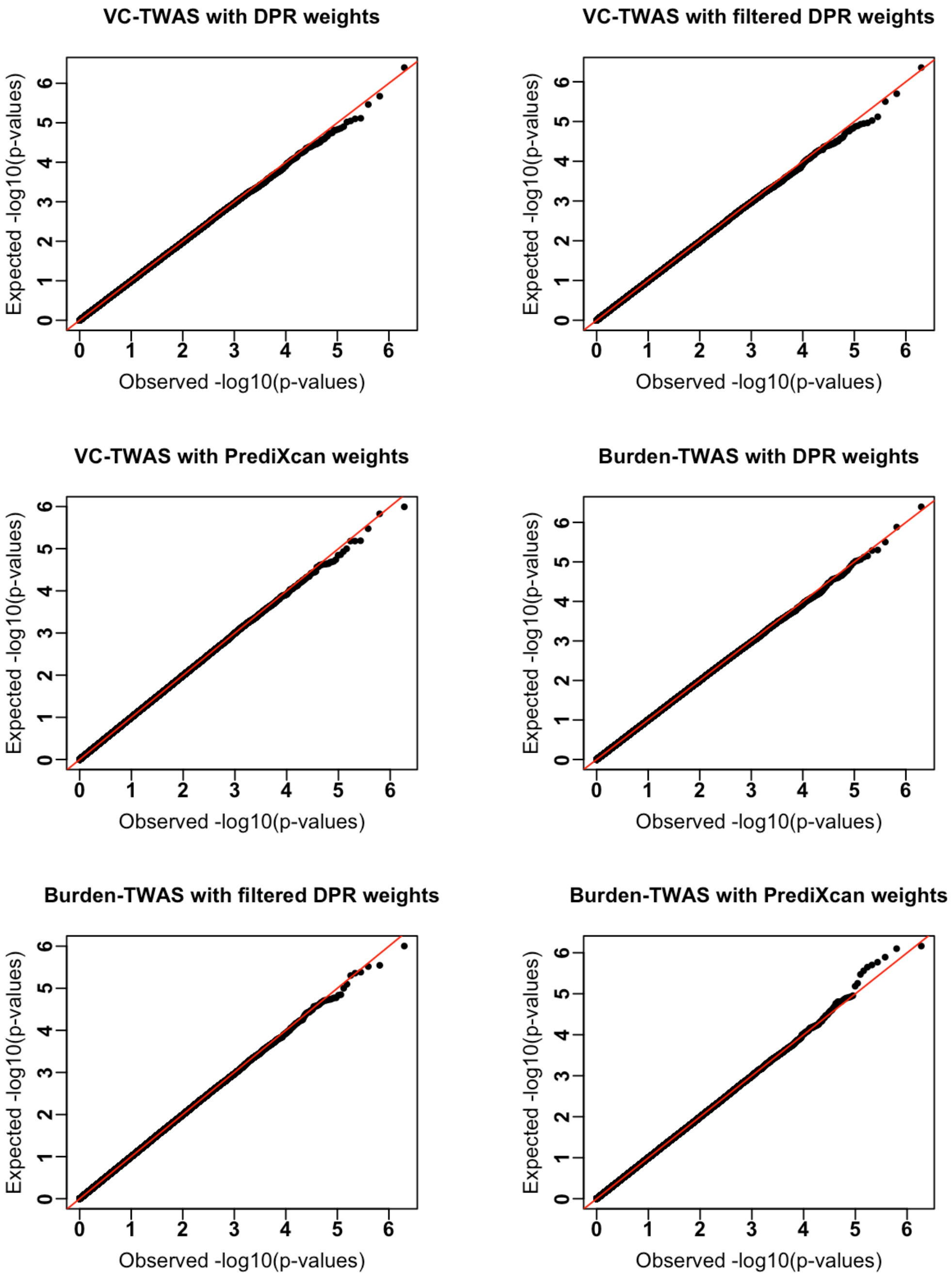


**Fig S4.** Q-Q plots for VC-TWAS and Burden-TWAS with DPR weights, filtered DPR weights, and PrediXcan weights under null hypothesis, where quantitative gene expression traits were generated with $p_{causal} = 0.2$ and $h_e^2$ = 0.1.

A)

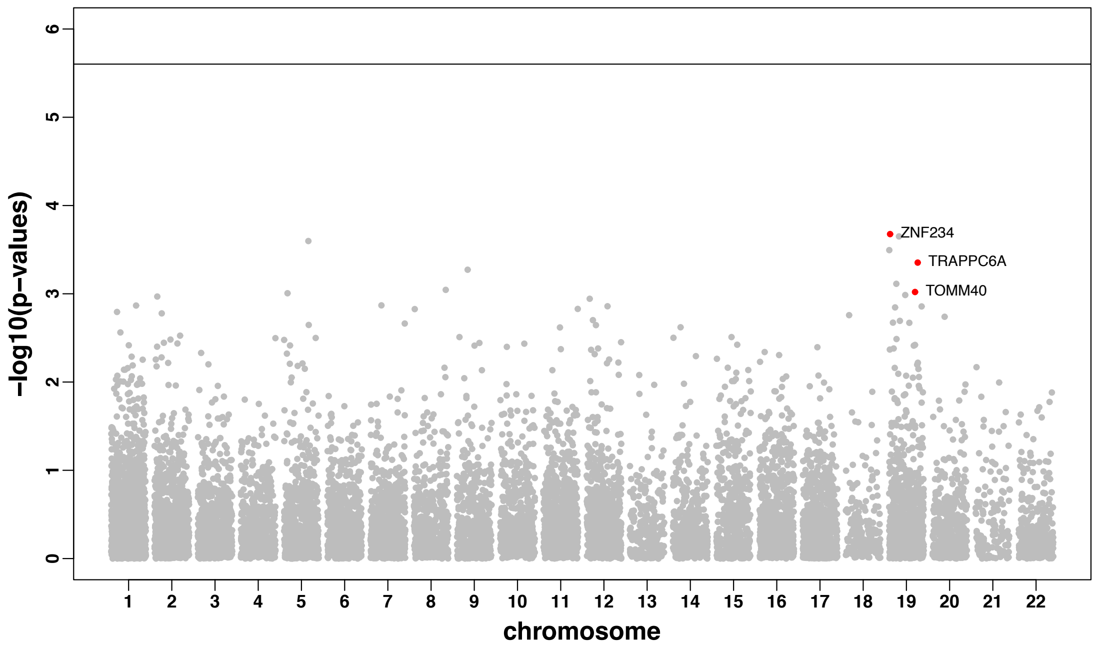


B)

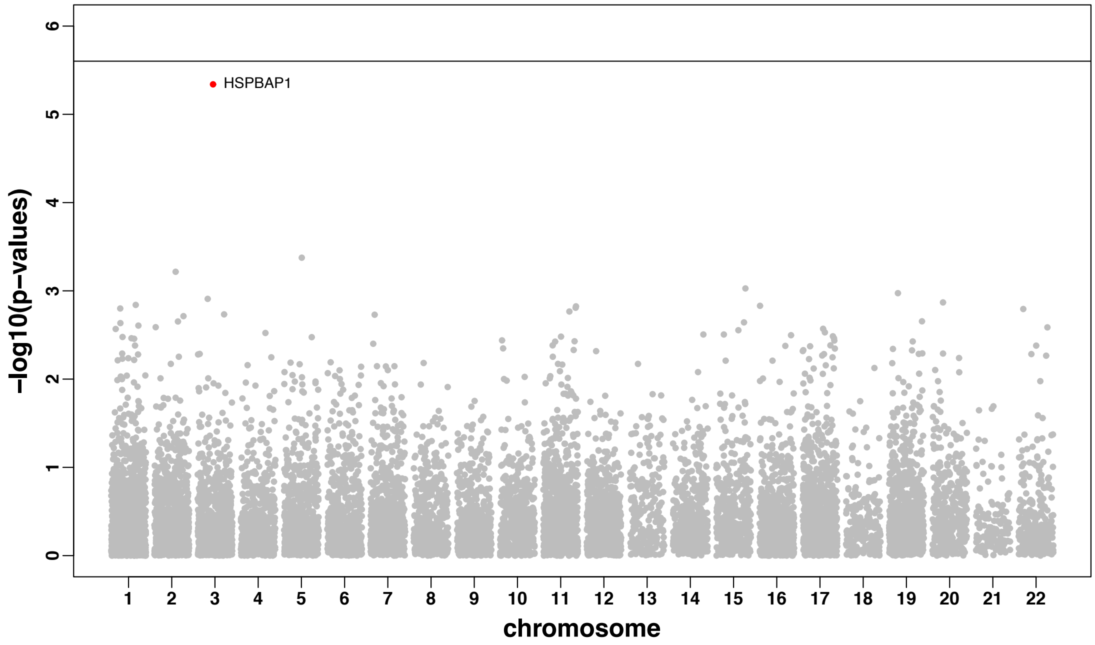


**Fig S5.** Manhattan plots of VC-TWAS results with filtered DPR weights for studying quantitative AD pathology of $\beta$-Amyloid (A) and tangles (B). Genes with FDA < 0.05 by meta VC-TWAS for studying AD clinical diagnosis are colored in red in (A) and top significant gene for studying tangles phenotype with FDR = 0.058 is colored in red in (B).

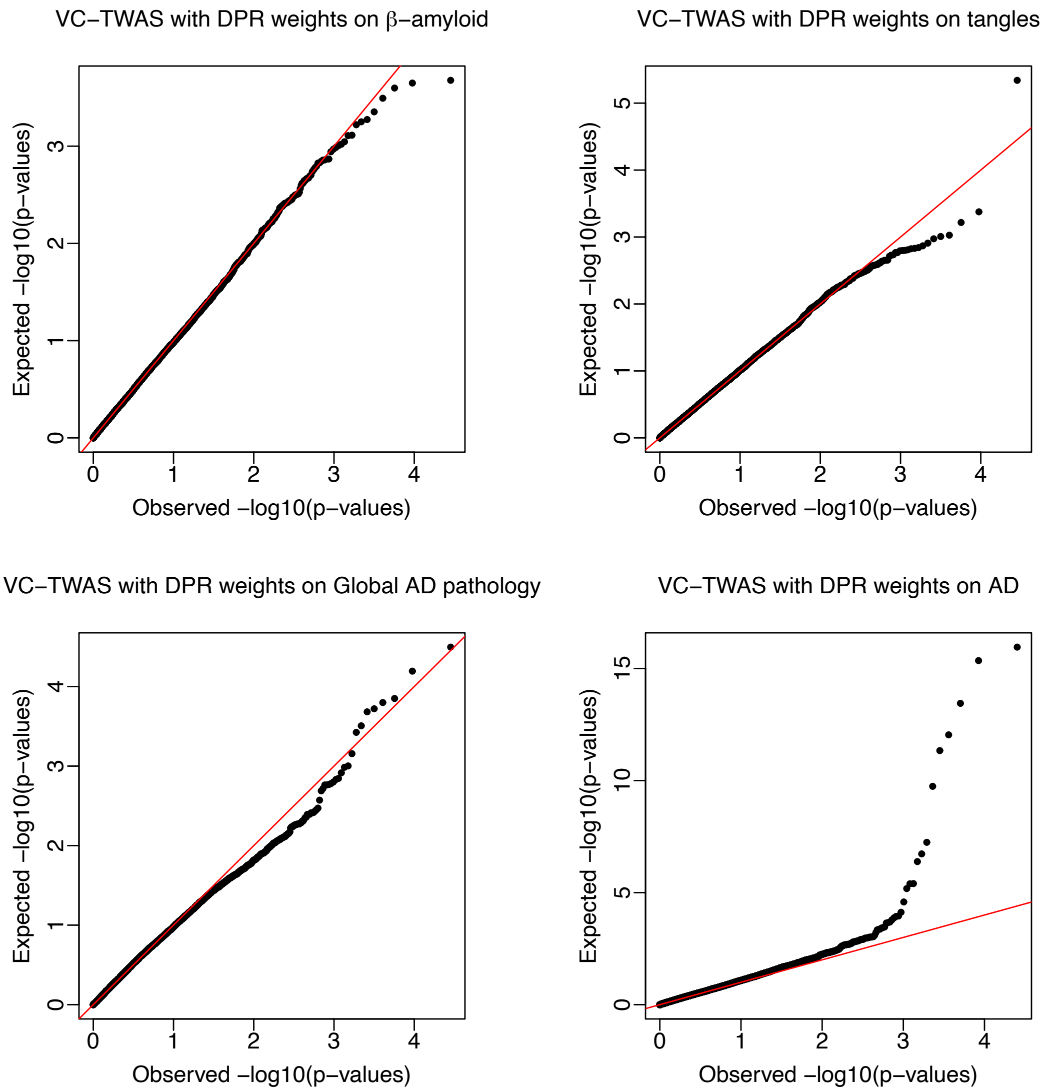


**Fig S6.** Q-Q plots of VC-TWAS results with filtered DPR weights for studying $\beta$-amyloid, tangles, and global AD pathology with ROS/MAP cohort, as well as meta VC-TWAS results with filtered DPR weights for studying AD clinical diagnosis with ROS/MAP and Mayo Clinic cohorts.

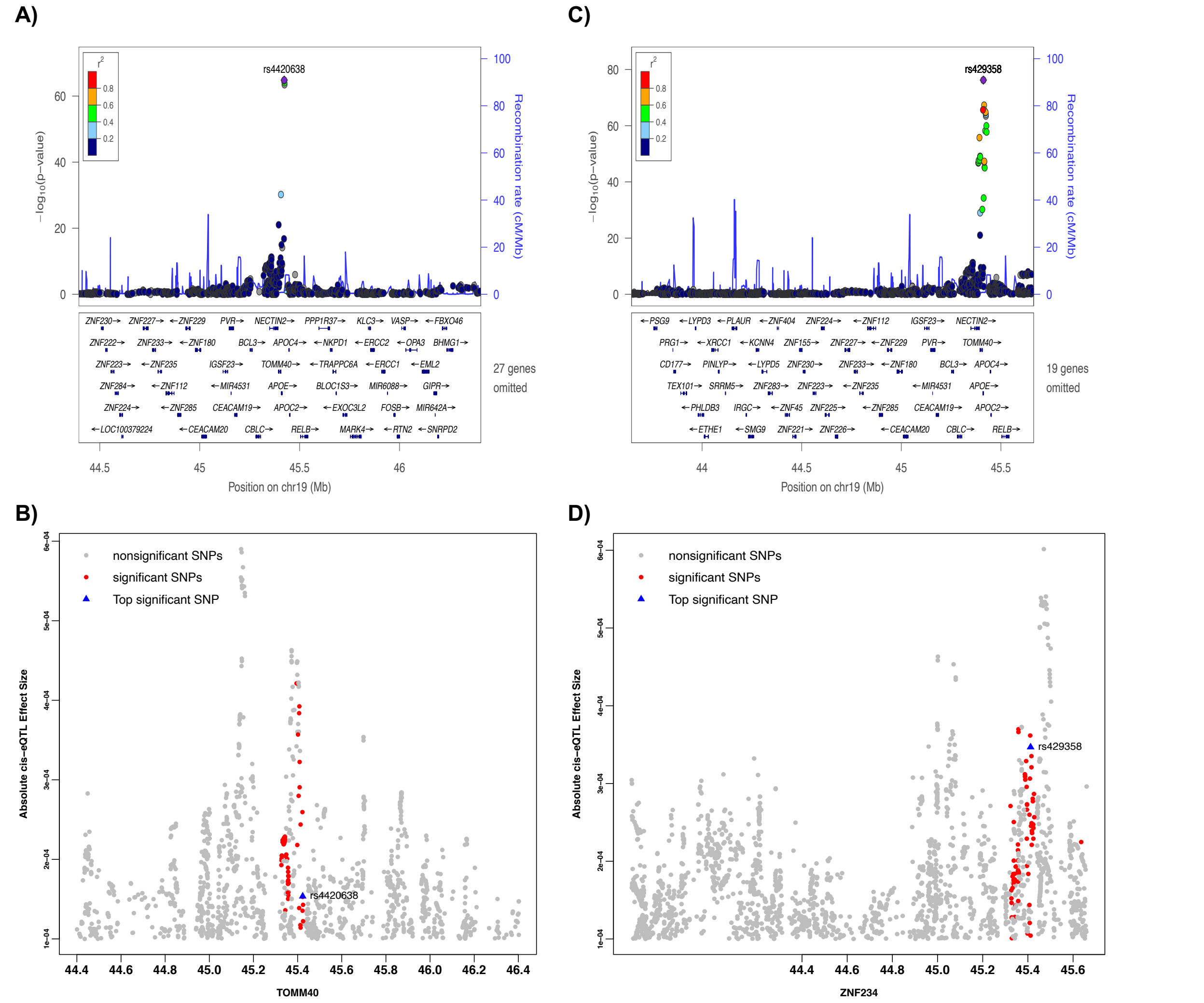


**Fig S7.** Locus zoom plots of GWAS results and the magnitude (i.e., absolute value) of cis-eQTL effect size estimates by DPR method for SNPs that were considered by VC-TWAS of genes *TOMM40* (A, B) and *ZNF2334* (C, D). Filtered test SNPs with the cis-eQTL effect size magnitude > $10^{-4}$ were plotted here. SNPs with GWAS p-value $< 5 \times 10^{-8}$ were colored in red in (B,D), top significant SNPs by GWAS in (A,C) were shown as the blue triangle in (B,D).

A)

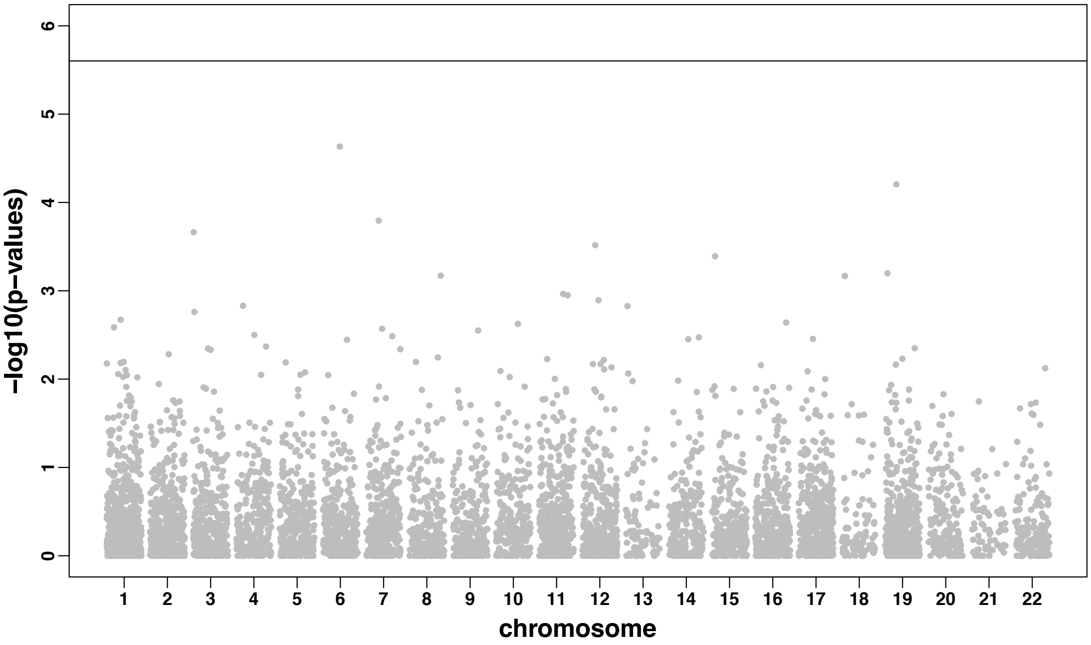


B)

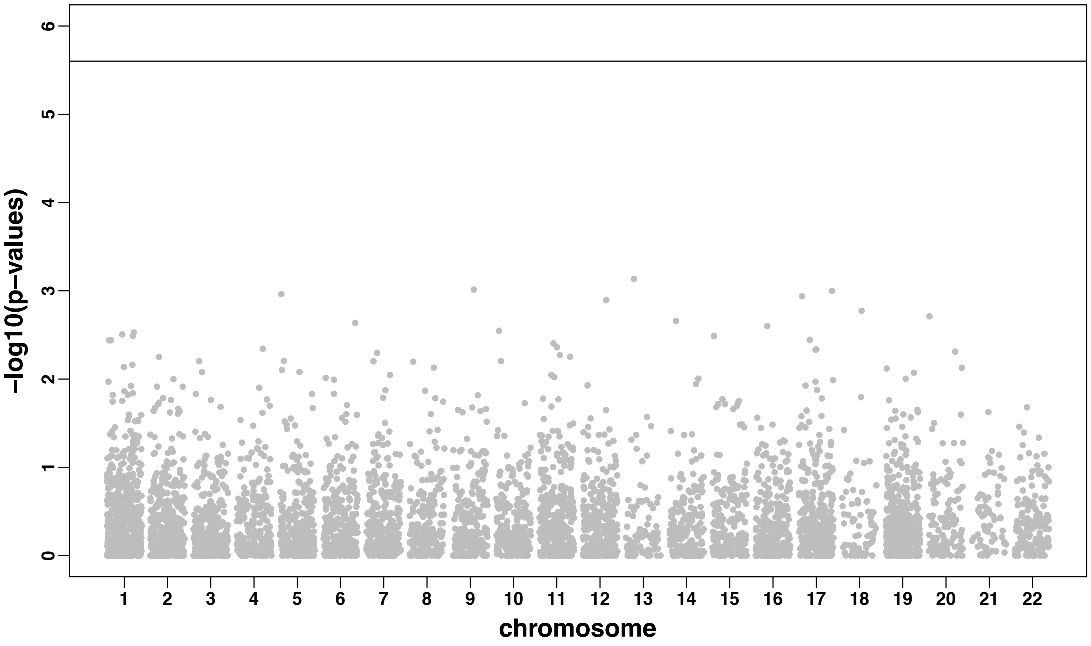


**Fig S8.** Manhattan plots of VC-TWAS results with PrediXcan weights for studying AD clinical diagnosis (A) and global AD pathology (B).

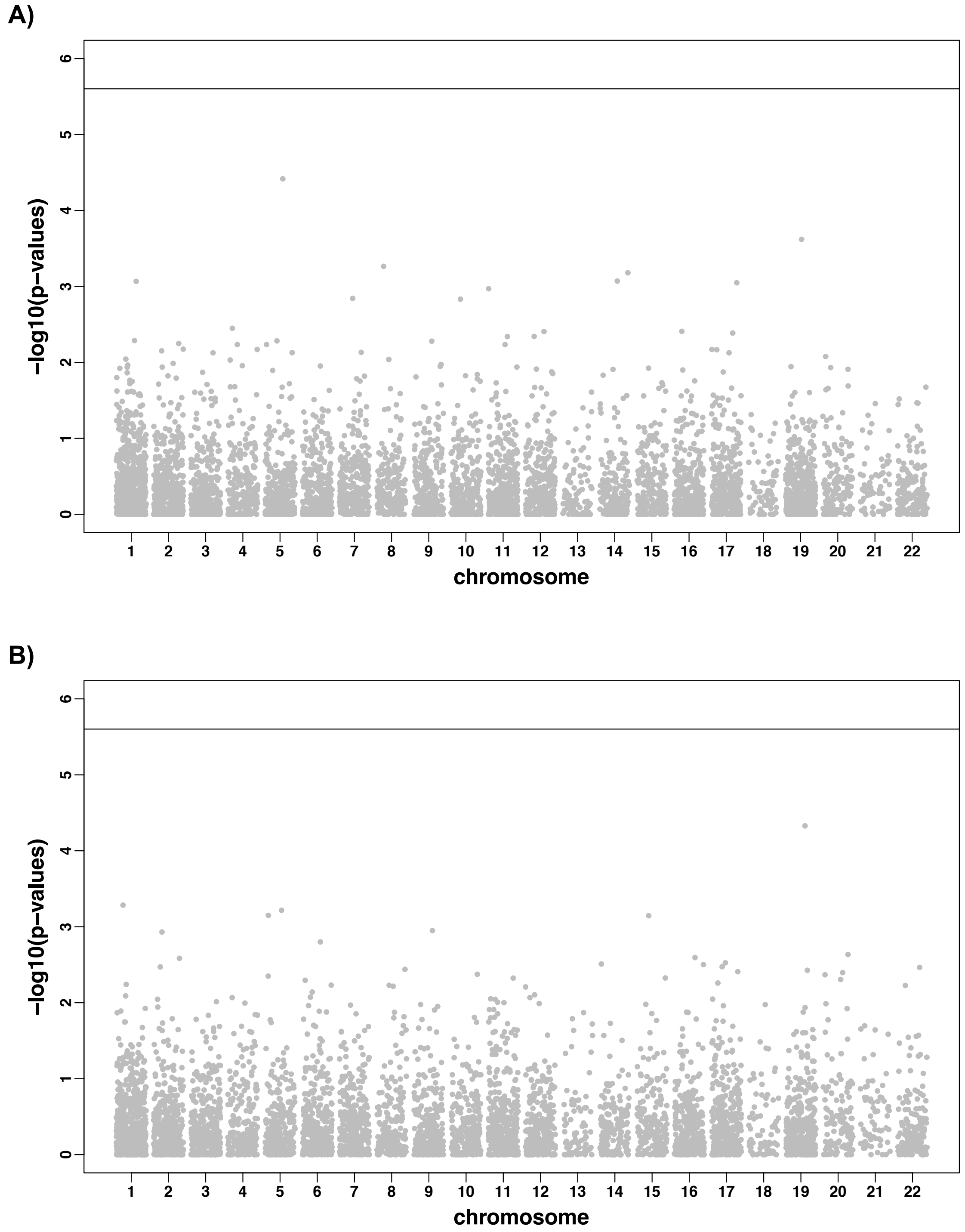


**Fig S9.** Manhattan plots of VC-TWAS results with PrediXcan weights for studying quantitative AD pathology of β-Amyloid (A) and tangles (B).

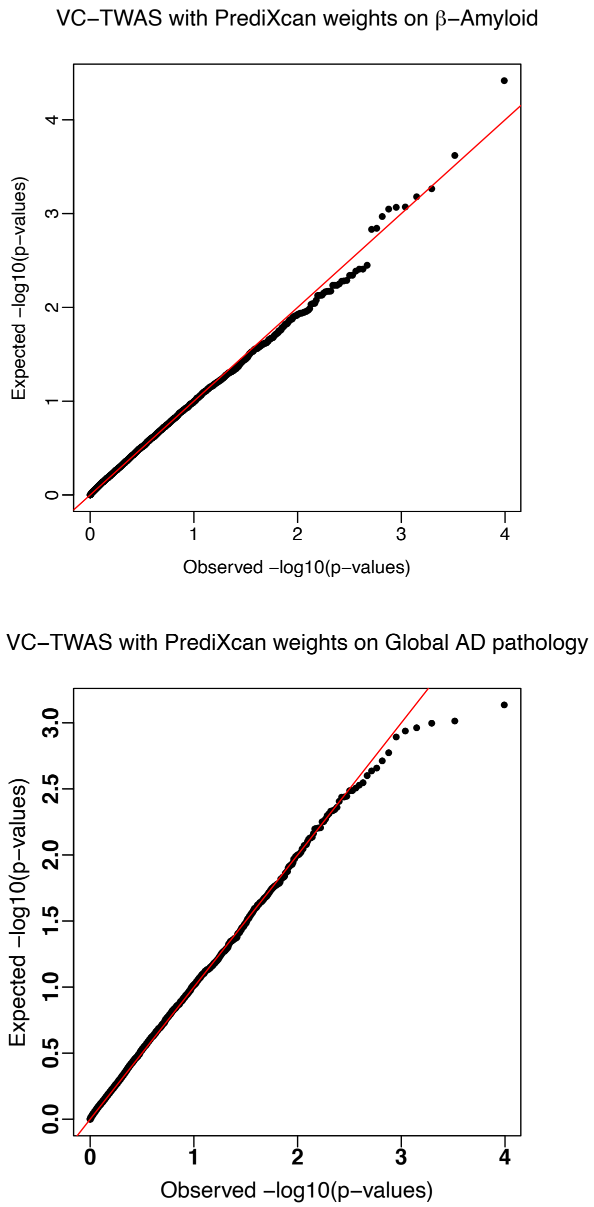


**Fig S10.** Q-Q plots of VC-TWAS results with PrediXcan weights for studying $\beta$-amyloid, tangles, and global AD pathology with ROS/MAO cohort, as well as meta VC-TWAS results with PrediXcan weights for studying AD clinical diagnosis with ROS/MAP and Mayo Clinic cohorts.

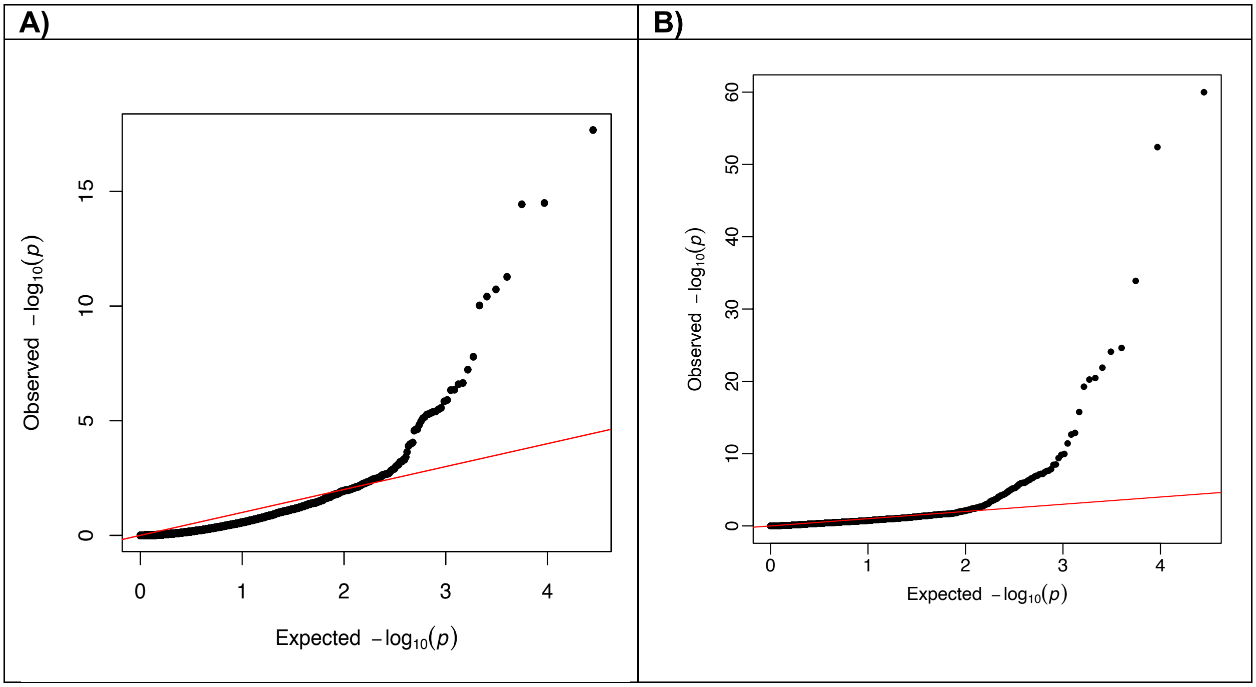


**Fig S11.** Q-Q plots of VC-TWAS results with cis-eQTL DPR filtered weights and BGW weights on IGAP GWAS summary statistics.

**Supplementary Tables**

| SNP number | Burden-TWAS | Burden-TWAS-SS | VC-TWAS | VC-TWAS-SS | CoMM | CoMM-SS |
|---|---|---|---|---|---|---|
| <1000 | 0.25 | 0.0016 | 0.28 | 0.15 | 73.62 | 2.11 |
| 1000-2000 | 0.28 | 0.0039 | 4.73 | 1.42 | 2333.62 | 81.93 |
| 2000-5000 | 0.26 | 0.0124 | 20.09 | 3.71 | 38037.01 | 373.16 |
| >5000 | 0.25 | 0.0478 | 138.93 | 20.64 | 83349.44 | 5476.43 |

**Table S1.** Computation time (in the unit of second) by Burden-TWAS, VC-TWAS and CoMM with individual and summary-level GWAS data by using with 1 CPU core with 32GB memory, for example genes that have different test SNP numbers.

| Gene | CHROM | $\beta$-Amyloid | Tangles | Global AD pathology |
|---|---|---|---|---|
| *ZNF234* | 19 | $2.10 \times 10^{-4*}$ | $1.06 \times 10^{-3*}$ | $6.39 \times 10^{-5*}$ |
| *CLASRP* | 19 | $1.39 \times 10^{-3}$ | $8.69 \times 10^{-3}$ | $3.76 \times 10^{-4*}$ |
| *TRAPPC6A* | 19 | $4.44 \times 10^{-4*}$ | $3.74 \times 10^{-3}$ | $1.91 \times 10^{-4*}$ |
| *TOMM40* | 19 | $9.55 \times 10^{-4*}$ | $6.95 \times 10^{-2}$ | $2.08 \times 10^{-4*}$ |
| *CEACAM19* | 19 | $1.03 \times 10^{-3*}$ | $1.21 \times 10^{-2}$ | $3.19 \times 10^{-5*}$ |

**Table S2.** Genes with VC-TWAS p-value <0.0013 with respect to at least one AD pathology phenotype and FDR <0.05 by meta VC-TWAS of AD clinical diagnosis. AD risk genes identified by previous GWAS are shaded in grey.

| Gene name | CHROM | Start | End | P-value | FDR |
|---|---|---|---|---|---|
| *CUTA* | 6 | 33,384,218 | 33,386,094 | $1.95 \times 10^{-5}$ | $5.96 \times 10^{-3}$ |
| *CLU* | 8 | 27,454,433 | 27,472,548 | $2.01 \times 10^{-5}$ | $6.13 \times 10^{-3}$ |
| *OSBP* | 11 | 59,341,870 | 59,383,617 | $8.67 \times 10^{-6}$ | $2.84 \times 10^{-3}$ |
| *STX3* | 11 | 59,480,928 | 59,573,354 | $2.82 \times 10^{-5}$ | $8.09 \times 10^{-3}$ |
| *PRPF19* | 11 | 60,658,201 | 60,674,060 | $2.93 \times 10^{-6}$ | $1.00 \times 10^{-3}$ |
| *TMEM109* | 11 | 60,681,345 | 60,690,915 | $3.66 \times 10^{-5}$ | $1.03 \times 10^{-2}$ |
| *TMEM132A* | 11 | 60,691,934 | 60,704,631 | $3.56 \times 10^{-6}$ | $1.19 \times 10^{-3}$ |
| *ME3* | 11 | 86,152,149 | 86,383,678 | $1.52 \times 10^{-5}$ | $4.76 \times 10^{-3}$ |
| *ZNF221* | 19 | 44,455,379 | 44,471,752 | $5.73 \times 10^{-5}$ | $1.55 \times 10^{-2}$ |
| *ZNF230* | 19 | 44,507,076 | 44,518,072 | $8.75 \times 10^{-20}$ | $6.48 \times 10^{-17}$ |
| *ZNF222* | 19 | 44,529,493 | 44,537,260 | $1.58 \times 10^{-11}$ | $6.16 \times 10^{-9}$ |
| *ZNF284* [b] | 19 | 44,576,296 | 44,591,623 | $6.39 \times 10^{-11}$ | $2.43 \times 10^{-8}$ |
| *ZNF225* [a] | 19 | 44,617,547 | 44,637,255 | $6.77 \times 10^{-8}$ | $2.51 \times 10^{-5}$ |
| *ZNF234* [a, b] | 19 | 44,645,709 | 44,664,462 | $3.38 \times 10^{-57}$ | $1.19 \times 10^{-53}$ |
| *ZNF226* | 19 | 44,669,214 | 44,681,836 | $7.69 \times 10^{-7}$ | $2.70 \times 10^{-4}$ |
| *ZNF227* [b] | 19 | 44,716,690 | 44,741,420 | $4.83 \times 10^{-18}$ | $3.24 \times 10^{-15}$ |
| *ZNF233* | 19 | 44,754,317 | 44,815,771 | $2.78 \times 10^{-5}$ | $8.09 \times 10^{-3}$ |
| *ZFP112* [b] | 19 | 44,830,705 | 44,905,774 | $7.16 \times 10^{-13}$ | $3.05 \times 10^{-10}$ |
| *PVR* [b] | 19 | 45,147,097 | 45,169,429 | $3.28 \times 10^{-14}$ | $1.65 \times 10^{-11}$ |
| *CEACAM19* [a,b] | 19 | 45,174,723 | 45,187,631 | $7.27 \times 10^{-27}$ | $7.30 \times 10^{-24}$ |
| *BCL3* [b] | 19 | 45,250,961 | 45,263,301 | $3.09 \times 10^{-17}$ | $1.89 \times 10^{-14}$ |
| *BCAM* | 19 | 45,312,337 | 45,324,677 | $3.35 \times 10^{-13}$ | $1.47 \times 10^{-10}$ |
| *PVRL2* | 19 | 45,349,392 | 45,392,485 | $7.07 \times 10^{-23}$ | $5.85 \times 10^{-20}$ |
| *TOMM40* [a,b] | 19 | 45,394,476 | 45,406,935 | $1.52 \times 10^{-69}$ | $7.13 \times 10^{-66}$ |
| *APOE* | 19 | 45,408,955 | 45,412,650 | $2.59 \times 10^{-13}$ | $1.22 \times 10^{-10}$ |
| *APOC1* | 19 | 45,417,920 | 45,422,606 | $7.83 \times 10^{-12}$ | $3.16 \times 10^{-9}$ |
| *CLPTM1* [a,b] | 19 | 45,457,847 | 45,496,598 | $1.48 \times 10^{-30}$ | $1.60 \times 10^{-27}$ |
| *RELB* [a] | 19 | 45,504,694 | 45,541,452 | $7.48 \times 10^{-41}$ | $1.50 \times 10^{-37}$ |
| *CLASRP* [a,b] | 19 | 45,542,297 | 45,574,214 | $1.91 \times 10^{-90}$ | $2.69 \times 10^{-86}$ |
| *ZNF296* [b] | 19 | 45,574,758 | 45,579,845 | $4.05 \times 10^{-16}$ | $2.28 \times 10^{-13}$ |
| *GEMIN7* | 19 | 45,582,529 | 45,594,782 | $1.56 \times 10^{-16}$ | $9.17 \times 10^{-14}$ |
| *PPP1R37* | 19 | 45,595,049 | 45,651,335 | $1.02 \times 10^{-17}$ | $6.49 \times 10^{-15}$ |
| *NKPD1* | 19 | 45,653,007 | 45,663,408 | $2.61 \times 10^{-19}$ | $1.84 \times 10^{-15}$ |
| *TRAPPC6A* [a,b] | 19 | 45,666,186 | 45,681,485 | $5.33 \times 10^{-51}$ | $1.50 \times 10^{-47}$ |
| *BLOC1S3* | 19 | 45,682,002 | 45,685,057 | $6.28 \times 10^{-5}$ | $1.67 \times 10^{-2}$ |
| *MARK4* [a,b] | 19 | 45,754,549 | 45,808,541 | $3.16 \times 10^{-71}$ | $2.22 \times 10^{-67}$ |
| *ERCC2* | 19 | 45,854,245 | 45,873,876 | $4.76 \times 10^{-5}$ | $1.31 \times 10^{-2}$ |
| *PPP1R13L* [a] | 19 | 45,882,891 | 45,909,607 | $1.25 \times 10^{-37}$ | $1.95 \times 10^{-34}$ |
| *CD3EAP* [b] | 19 | 45,909,466 | 45,914,024 | $1.15 \times 10^{-7}$ | $4.15 \times 10^{-5}$ |
| *ERCC1* | 19 | 45,910,590 | 45,982,086 | $6.89 \times 10^{-14}$ | $3.34 \times 10^{-11}$ |
| *FOSB* | 19 | 45,971,252 | 45,978,414 | $4.06 \times 10^{-21}$ | $3.17 \times 10^{-18}$ |
| *RTN2* [b] | 19 | 45,988,549 | 46,000,313 | $1.89 \times 10^{-26}$ | $1.78 \times 10^{-23}$ |
| *PPM1N* [b] | 19 | 45,992,034 | 46,005,768 | $1.30 \times 10^{-15}$ | $7.04 \times 10^{-13}$ |
| *VASP* | 19 | 46,010,687 | 46,030,236 | $3.11 \times 10^{-41}$ | $7.28 \times 10^{-38}$ |

| | | | | | |
|---|---|---|---|---|---|
| *GPR4* | 19 | 46,093,024 | 46,105,466 | $8.88 \times 10^{-6}$ | $2.84 \times 10^{-3}$ |
| *EML2* [a,b] | 19 | 46,112,659 | 46,148,726 | $1.83 \times 10^{-34}$ | $2.35 \times 10^{-31}$ |
| *GIPR* [a,b] | 19 | 46,171,501 | 46,185,704 | $4.21 \times 10^{-25}$ | $3.70 \times 10^{-22}$ |
| *SNRPD2* [b] | 19 | 46,190,712 | 46,195,443 | $9.97 \times 10^{-15}$ | $5.19 \times 10^{-12}$ |
| *QPCTL* | 19 | 46,195,740 | 46,207,240 | $7.87 \times 10^{-12}$ | $3.16 \times 10^{-9}$ |
| *FBXO46* [a,b] | 19 | 46,213,886 | 46,234,151 | $1.74 \times 10^{-36}$ | $2.45 \times 10^{-33}$ |
| *DMPK* | 19 | 46,272,977 | 46,285,815 | $6.79 \times 10^{-32}$ | $7.95 \times 10^{-29}$ |
| *DMWD* | 19 | 46,286,204 | 46,296,060 | $1.40 \times 10^{-4}$ | $3.57 \times 10^{-2}$ |
| *SYMPK* | 19 | 46,318,692 | 46,366,548 | $1.54 \times 10^{-4}$ | $3.87 \times 10^{-2}$ |
| *IRF2BP1* [b] | 19 | 46,386,865 | 46,389,376 | $5.02 \times 10^{-39}$ | $8.83 \times 10^{-36}$ |
| *MYPOP* [b] | 19 | 46,393,284 | 46,405,862 | $2.80 \times 10^{-13}$ | $1.27 \times 10^{-10}$ |
| *NLRP2* | 19 | 55,476,437 | 55,512,510 | $2.82 \times 10^{-5}$ | $8.09 \times 10^{-3}$ |
| *HAR1A* | 20 | 61,733,556 | 61,735,738 | $1.17 \times 10^{-4}$ | $3.05 \times 10^{-2}$ |

a. Genes also identified as significant by VC-TWAS with individual-level GWAS data of ROS/MAP and Mayo Clinic cohorts.
b. Genes identified as significant by both VC-TWAS and Burden-TWAS using IGAP summary statistics with filtered cis-eQTL DPR weights.

**Table S3.** Significant genes identified by VC-TWAS using IGAP summary statistics with filtered cis-eQTL DPR weights. Significant genes were identified with FDR < 0.05. AD risk genes identified by previous GWAS are shaded in grey.

| Gene Name | CHROM | Start | End | P-value | FDR |
|---|---|---|---|---|---|
| PPIEL | 1 | 39,997,509 | 40,024,379 | $1.41 \times 10^{-4}$ | $3.01 \times 10^{-2}$ |
| ARHGAP29 | 1 | 94,614,543 | 94,740,624 | $1.14 \times 10^{-7}$ | $6.11 \times 10^{-5}$ |
| RWDD3 | 1 | 95,699,710 | 95,712,781 | $4.17 \times 10^{-53}$ | $2.94 \times 10^{-49}$ |
| GPATCH2 | 1 | 217,600,333 | 217,804,424 | $2.41 \times 10^{-7}$ | $1.17 \times 10^{-4}$ |
| RP11-211A18.2 | 1 | 227,421,969 | 227,423,056 | $4.84 \times 10^{-6}$ | $1.59 \times 10^{-3}$ |
| C2orf50 | 2 | 11,273,178 | 11,286,916 | $4.92 \times 10^{-5}$ | $1.20 \times 10^{-2}$ |
| ANKRD30BL | 2 | 132,905,163 | 133,015,542 | $1.43 \times 10^{-8}$ | $1.06 \times 10^{-5}$ |
| TTC21B | 2 | 166,713,984 | 166,810,353 | $4.86 \times 10^{-7}$ | $2.14 \times 10^{-4}$ |
| SPEG | 2 | 220,299,567 | 220,363,009 | $1.11 \times 10^{-10}$ | $1.12 \times 10^{-7}$ |
| FBXO36 | 2 | 230,787,017 | 230,877,825 | $3.14 \times 10^{-9}$ | $2.60 \times 10^{-6}$ |
| PTH1R | 3 | 46,919,235 | 46,945,287 | $1.52 \times 10^{-6}$ | $5.50 \times 10^{-4}$ |
| ZXDC | 3 | 126,156,443 | 126,194,762 | $7.11 \times 10^{-6}$ | $2.13 \times 10^{-3}$ |
| XPO5 | 6 | 43,490,071 | 43,543,812 | $3.76 \times 10^{-8}$ | $2.40 \times 10^{-5}$ |
| RP1-180E22.3 | 6 | 52,442,282 | 52,444,325 | $2.14 \times 10^{-4}$ | $4.30 \times 10^{-2}$ |
| BCLAF1 | 6 | 136,578,000 | 136,610,989 | $6.31 \times 10^{-7}$ | $2.69 \times 10^{-4}$ |
| AC017116.8 | 7 | 44,078,769 | 44,081,905 | $6.18 \times 10^{-8}$ | $3.70 \times 10^{-5}$ |
| MRPL15 | 8 | 55,047,769 | 55,060,461 | $6.21 \times 10^{-5}$ | $1.43 \times 10^{-2}$ |
| VCPIP1 | 8 | 67,540,721 | 67,579,452 | $3.52 \times 10^{-9}$ | $2.75 \times 10^{-6}$ |
| GEM | 8 | 95,261,480 | 95,274,578 | $5.51 \times 10^{-5}$ | $1.31 \times 10^{-2}$ |
| *TRAF1* | 9 | 123,664,670 | 123,691,451 | $5.69 \times 10^{-21}$ | $1.00 \times 10^{-17}$ |
| PBX3 | 9 | 128,509,623 | 128,729,656 | $1.23 \times 10^{-6}$ | $4.56 \times 10^{-4}$ |
| C9orf167 | 9 | 140,172,200 | 140,177,093 | $5.66 \times 10^{-6}$ | $1.81 \times 10^{-3}$ |
| AGAP10 | 10 | 47,191,843 | 47,239,738 | $1.29 \times 10^{-5}$ | $3.64 \times 10^{-3}$ |
| HELLS | 10 | 96,305,546 | 96,373,662 | $3.51 \times 10^{-21}$ | $7.05 \times 10^{-18}$ |
| PKD2L1 | 10 | 102,047,902 | 102,090,243 | $1.03 \times 10^{-60}$ | $1.45 \times 10^{-56}$ |
| C10orf137 | 10 | 127,408,083 | 127,452,712 | $3.26 \times 10^{-6}$ | $1.09 \times 10^{-3}$ |
| SLC39A13 | 11 | 47,428,682 | 47,438,047 | $1.81 \times 10^{-4}$ | $3.76 \times 10^{-2}$ |
| LETMD1 | 12 | 51,441,744 | 51,454,207 | $6.19 \times 10^{-6}$ | $1.94 \times 10^{-3}$ |
| ZBTB39 | 12 | 57,392,617 | 57,400,230 | $8.73 \times 10^{-5}$ | $1.95 \times 10^{-2}$ |
| KITLG | 12 | 88,885,884 | 88,974,628 | $1.74 \times 10^{-4}$ | $3.66 \times 10^{-2}$ |
| NT5DC3 | 12 | 104,164,230 | 104,234,975 | $3.95 \times 10^{-12}$ | $4.28 \times 10^{-9}$ |
| SERTM1 | 13 | 37,248,048 | 37,271,976 | $1.97 \times 10^{-4}$ | $4.03 \times 10^{-2}$ |
| SEC23A | 14 | 39,501,122 | 39,578,850 | $8.85 \times 10^{-5}$ | $1.95 \times 10^{-2}$ |
| KTN1-AS1 | 14 | 55,965,995 | 56,046,828 | $1.09 \times 10^{-6}$ | $4.25 \times 10^{-4}$ |
| ACOT4 | 14 | 74,058,409 | 74,063,200 | $1.30 \times 10^{-34}$ | $6.10 \times 10^{-31}$ |
| RP13-487P22.1 | 15 | 25,590,779 | 25,592,382 | $3.82 \times 10^{-5}$ | $9.60 \times 10^{-3}$ |
| C15orf58 | 15 | 90,777,039 | 90,785,315 | $1.17 \times 10^{-7}$ | $6.11 \times 10^{-5}$ |

| | | | | | |
|---|---|---|---|---|---|
| IGSF6 | 16 | 21,652,608 | 21,663,981 | $1.37 \times 10^{-13}$ | $1.75 \times 10^{-10}$ |
| AC012146.7 | 17 | 5,014,762 | 5,018,299 | $5.58 \times 10^{-20}$ | $8.74 \times 10^{-17}$ |
| AC092296.1 | 19 | 36,804,643 | 36,822,602 | $2.24 \times 10^{-13}$ | $2.63 \times 10^{-10}$ |
| GNAS | 20 | 57,414,772 | 57,486,247 | $8.07 \times 10^{-25}$ | $2.27 \times 10^{-21}$ |
| FOXRED2 | 22 | 36,883,236 | 36,903,148 | $1.05 \times 10^{-6}$ | $4.24 \times 10^{-4}$ |
| H1F0 | 22 | 38,201,113 | 38,203,442 | $8.26 \times 10^{-5}$ | $1.88 \times 10^{-2}$ |
| PPPDE2 | 22 | 41,994,031 | 42,017,100 | $4.15 \times 10^{-10}$ | $3.65 \times 10^{-7}$ |
| PPP6R2 | 22 | 50,781,732 | 50,883,514 | $6.97 \times 10^{-21}$ | $4.77 \times 10^{-2}$ |

**Table S4:** Novel significant genes identified by VC-TWAS using summary statistics with BGW weights on IGAP summary statistics.
